## Supplementary Information for "The catalytic mechanism of the RNA methyltransferase METTL3"

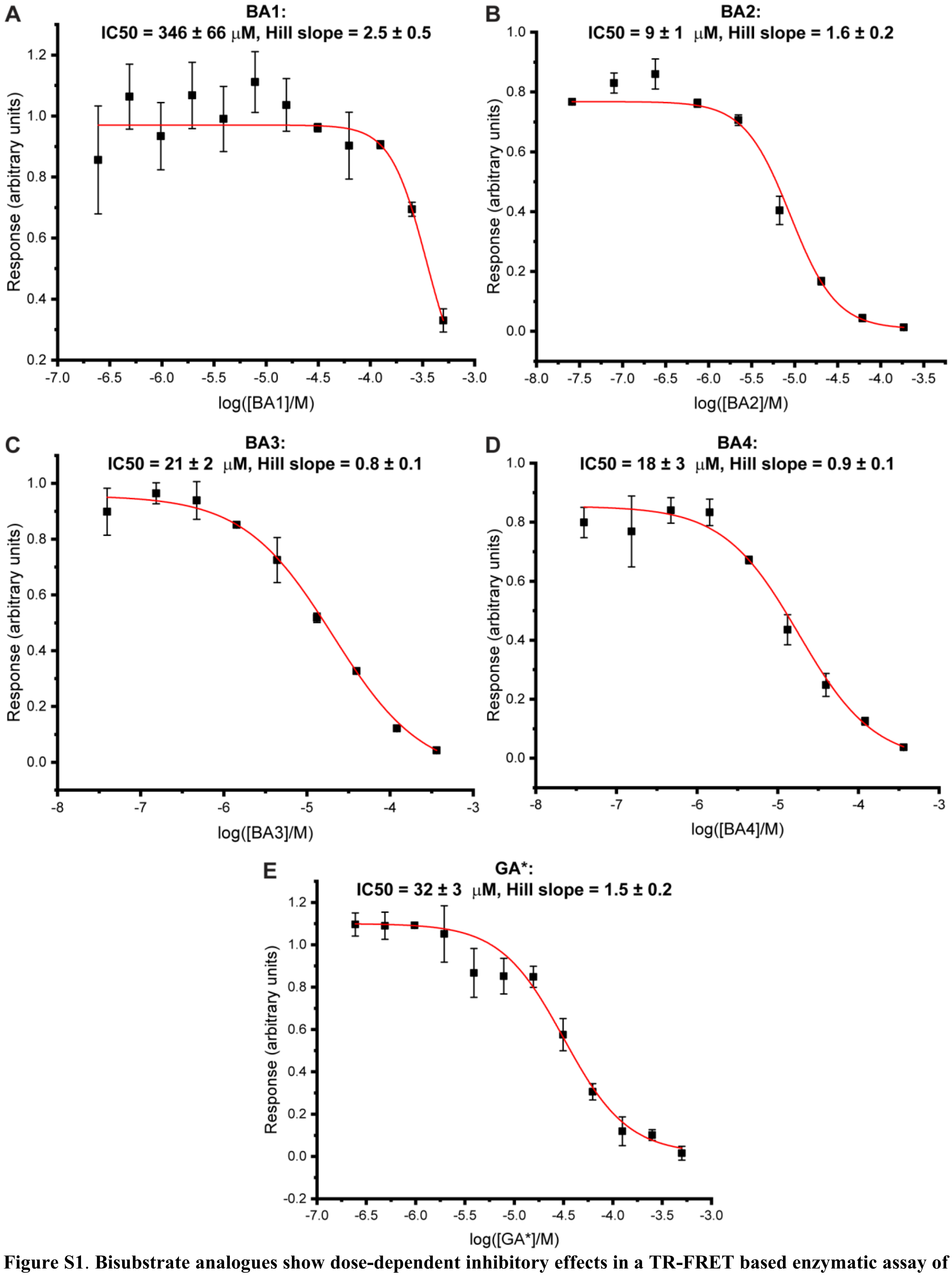


**Figure S1**. **Bisubstrate analogues show dose-dependent inhibitory effects in a TR-FRET based enzymatic assay of METTL3-14**. Dose-response curves derived from the reader-based TR-FRET inhibition assay on METTL3-14 (mean ± SD, n = 2 or 3 technical replicates) for BAs for which $IC_{50}$ values could be determined: BA1 (**A**), BA2 (**B**), BA3 (**C**), BA4 (**D**), and GA* (**E**). $IC_{50}$ and Hill Slope values were obtained from fits with nonlinear regression “log(inhibitor) vs. normalized response with variable slope” and are give at the top of each curve.

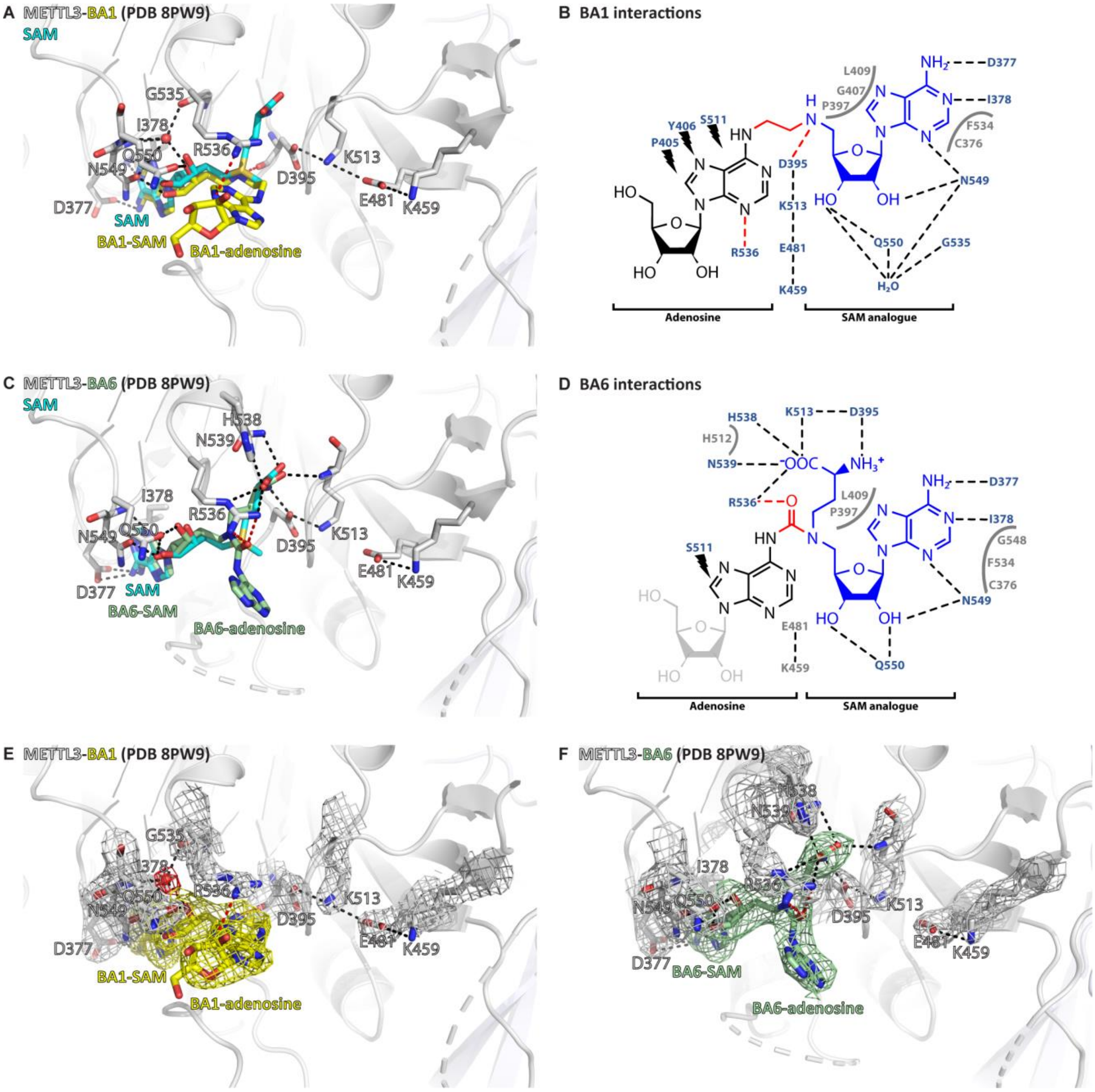


**Figure S2**. **Crystal structures of the complex of METTL3-14 with the bisubstrate analogues BA1 and BA6 show divergent interactions with the SAM moiety**. (**A**) Structure of the METTL3-BA1 complex. METTL3 backbone is shown as ribbon, sidechains involved in polar interactions with BA1 or intramolecularly are shown as sticks. Waters are shown as red spheres. BA1 (yellow) and SAM (cyan) are shown as sticks, SAM and the BA1 moieties are indicated. Because of the missing methionine moiety in BA1, METTL3 residues D395 and R536 cannot form natural salt bridges with the $NH_3^+$ and $COO^-$ groups of the methionine group of the SAM moiety, respectively, and instead form artificial hydrogen bonds with the BA linker and adenine ring, respectively (red dashes). (**B**) Ligplot+ analysis of the interaction between METTL3 and BA1. The SAM analogue and adenosine parts of the BA are indicated. Black dashed lines indicate polar contacts between METTL3 and BA1 in the crystal structure. Small lightnings highlight residues in METTL3 involved in hydrophobic contacts with the adenosine part of the BA. Residues forming the binding pocket environment are shown in grey. Red dashed lines indicate polar contacts that can only form because the methionine moiety of the SAM part of the BA is missing. (**C**) Structure of the METTL3-BA6 complex. METTL3 backbone is shown as ribbon, sidechains involved in polar interactions with BA6 or intramolecularly are shown as sticks. Waters are shown as red spheres. BA6 (palegreen) and SAM (cyan) are shown as sticks, SAM and the BA6 moieties are indicated. Note that BA6 is missing the ribose moiety of the substrate adenosine part due to lack of electron density in the crystal structure, probably due to flexibility of this group. Because of the polar urea group in the BA6 linker, METTL3 residue R536 forms a hydrogen bond with it (red dashes) leading to a shift of the position of the SAM-like moiety of BA6 compared to the natural SAM cosubstrate. (**D**) Ligplot+ analysis of the interaction between METTL3 and BA6, as in (B). The missing ribose of the substrate adenosine moiety in the crystal structure is indicated with a lighter colour. (**E,F**) Structures from (A) and (C) shown with electron densities for the BAs (contoured at 0.7 sigma) and METTL3 sidechains and waters (contoured at 1.0 sigma).

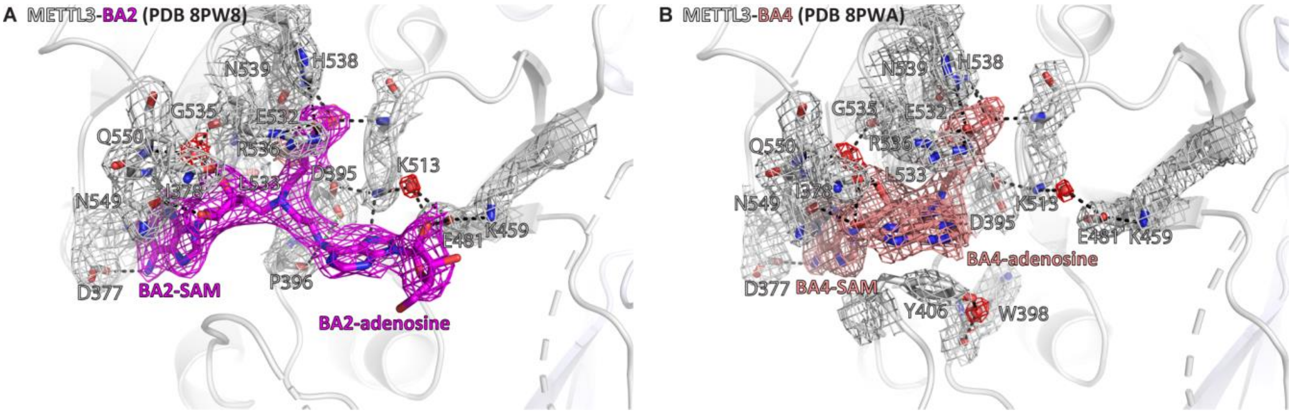


**Figure S3**. **Crystal structures of the complex of METTL3-14 with BA2 and BA4 show electron density supporting the conformations of the BAs and their interactions with METTL3**. Structures of METTL3 bound to BA2 (**A**) and BA4 (**B**) from main text Figure 3C and E, respectively, shown with electron densities for the BAs (contoured at 0.7 sigma) and METTL3 sidechains and waters (contoured at 1.0 sigma). METTL3 backbone is shown as ribbon, sidechains involved in polar contacts with the BAs or intramolecularly are shown as sticks, waters as red spheres. Dashes indicate polar contacts. The SAM and adenosine moieties of the BAs are indicated. Note that BA4 is missing electron density for the ribose of the substrate adenosine moiety probably due to flexibility of this group.

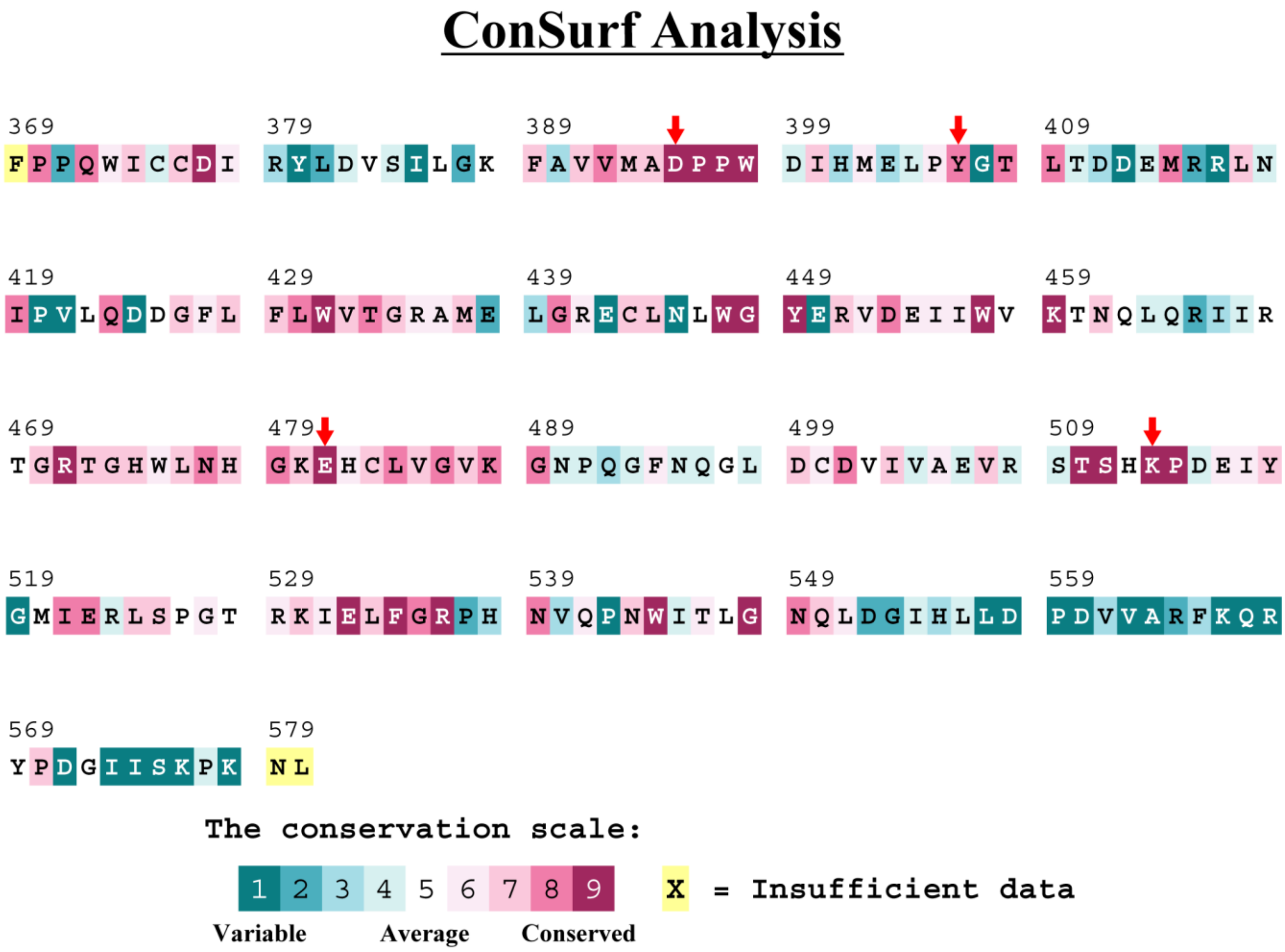


**Figure S4**. **METTL3 residues that interact with the adenosine part of the bisubstrate analogues are highly conserved**. Conservation analysis of the METTL3 MTase domain from PDB ID 5IL0 using Consurf-DB.[1,2] Relative conservation scores indicated by colours. The mutated residues in this study are marked with red arrows.

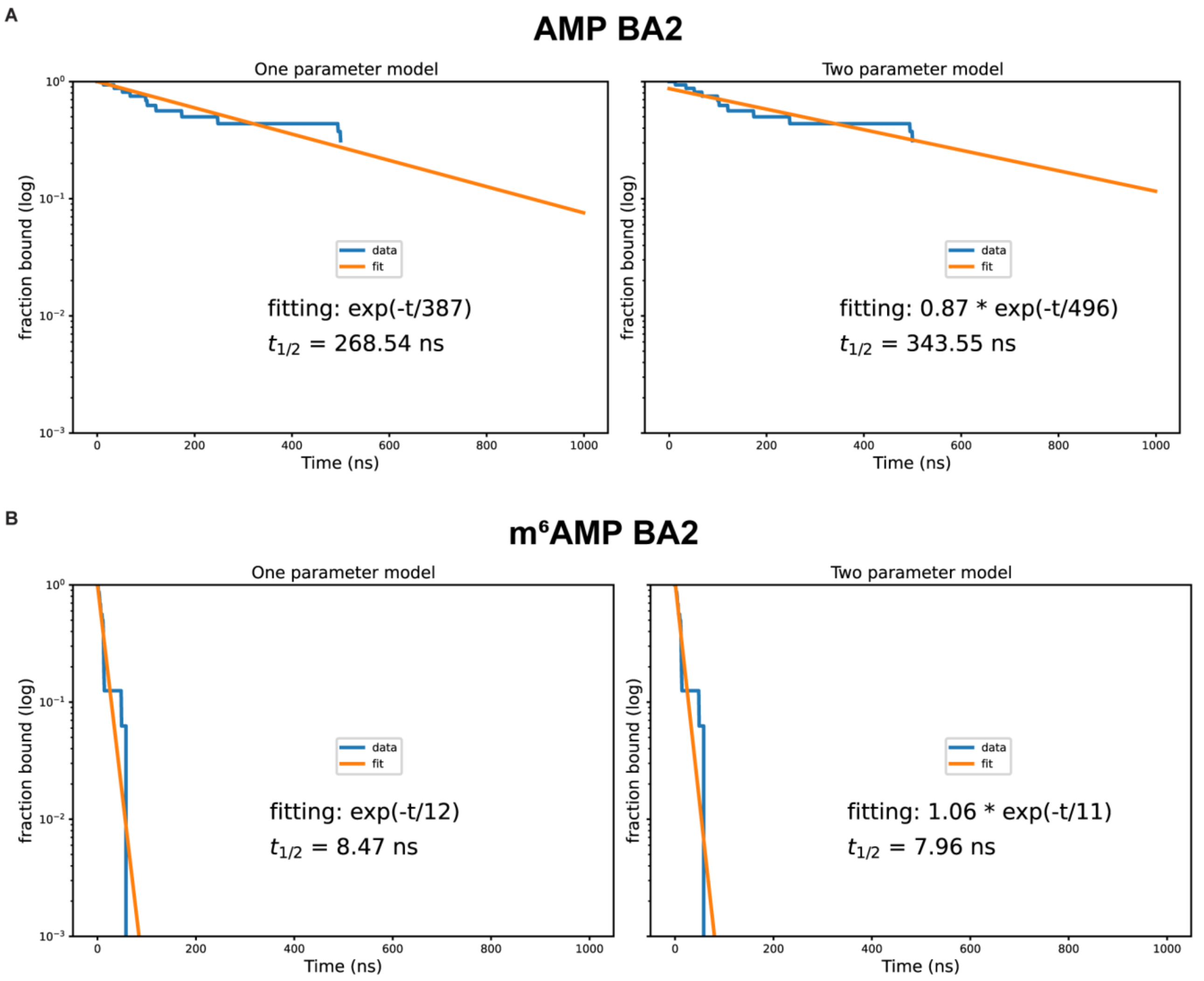


**Figure S5. Exponential fitting of AMP and m$^6$AMP dissociation.** Fitting done for AMP (**A**) and m$^6$AMP (**B**) with a one-parameter exponential decay function (left panel) and a two-parameter exponential decay function with a multiplicative factor (right panel). Starting conformation and bound ligand indicated at top of each Figure. Fitting done using all trajectories in contrast with values in Table 2.

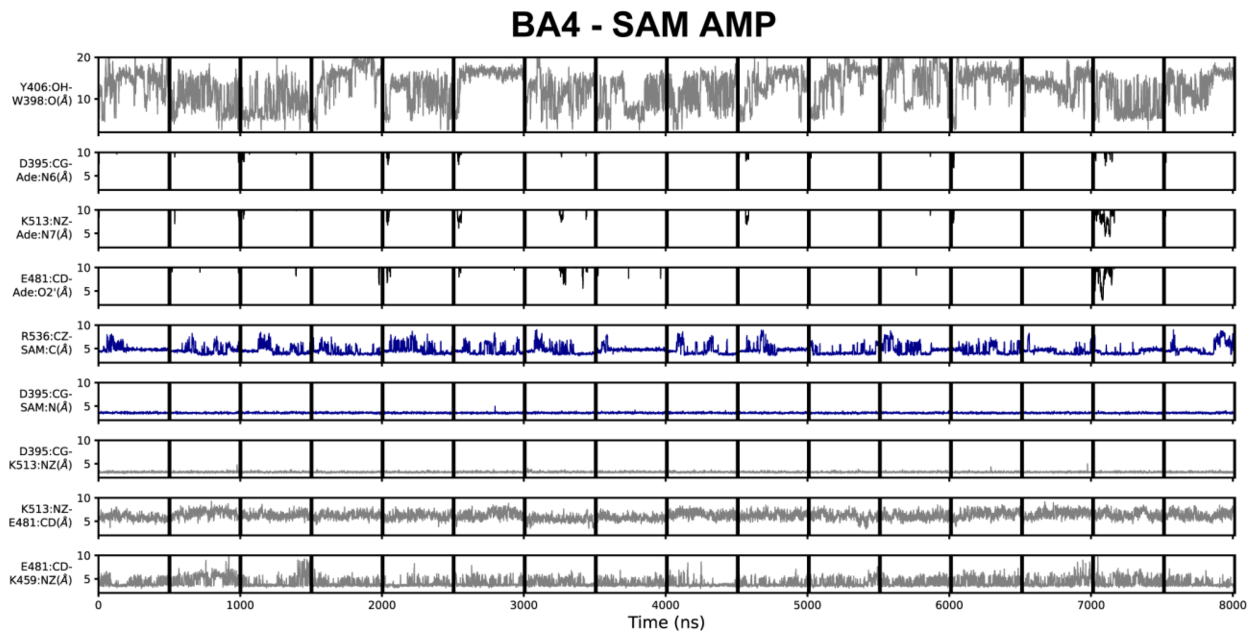


**Figure S6. Geometric annotation for trajectories started with substrates in the BA4 crystal structure conformation**. Shown are distance time series of 500-ns MD trajectories started from the BA4 conformation of METTL3 with (co)substrates. Starting conformation and bound ligands indicated on top of the Figure. Y406 to W398 distance, interaction of METTL3 to AMP substrate (black traces), METTL3 to SAM (blue traces), and intramolecular salt bridges.

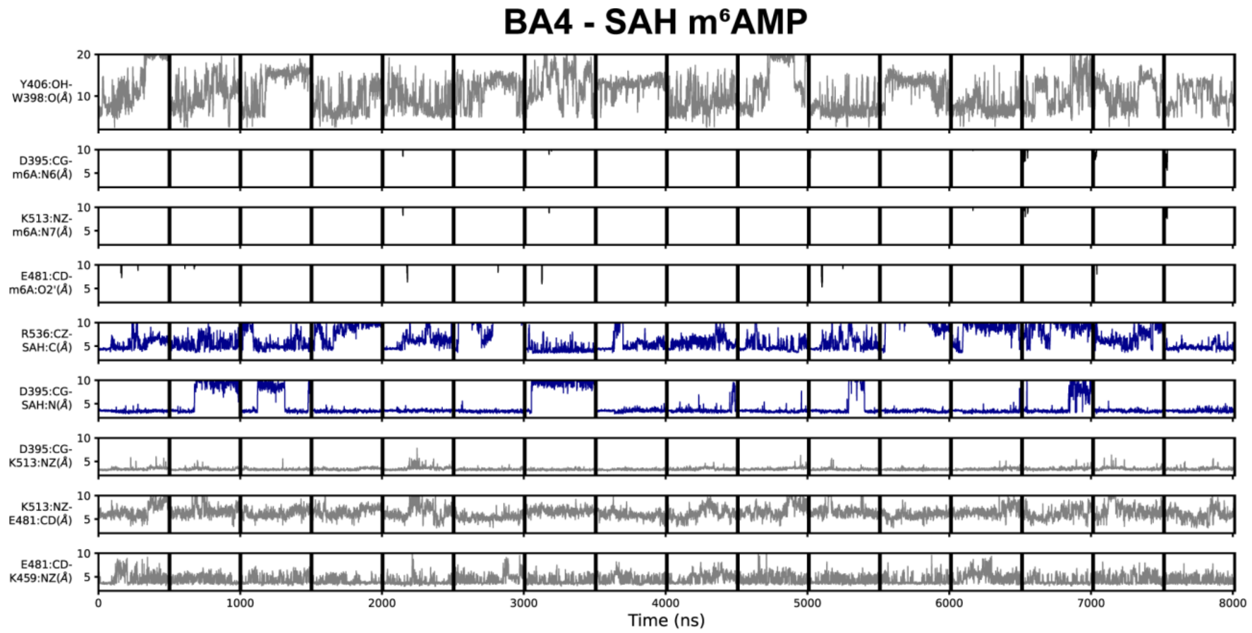


**Figure S7. Geometric annotation for trajectories started with products in the BA4 crystal structure conformation**. Shown are distance time series of 500-ns MD trajectories started from the BA4 conformation of METTL3 with (co)products. Starting conformation and bound ligands indicated on top of the Figure. Y406 to W398 distance, interaction of METTL3 to m$^6$AMP substrate (black traces), METTL3 to SAH (blue traces), and intramolecular salt bridges.

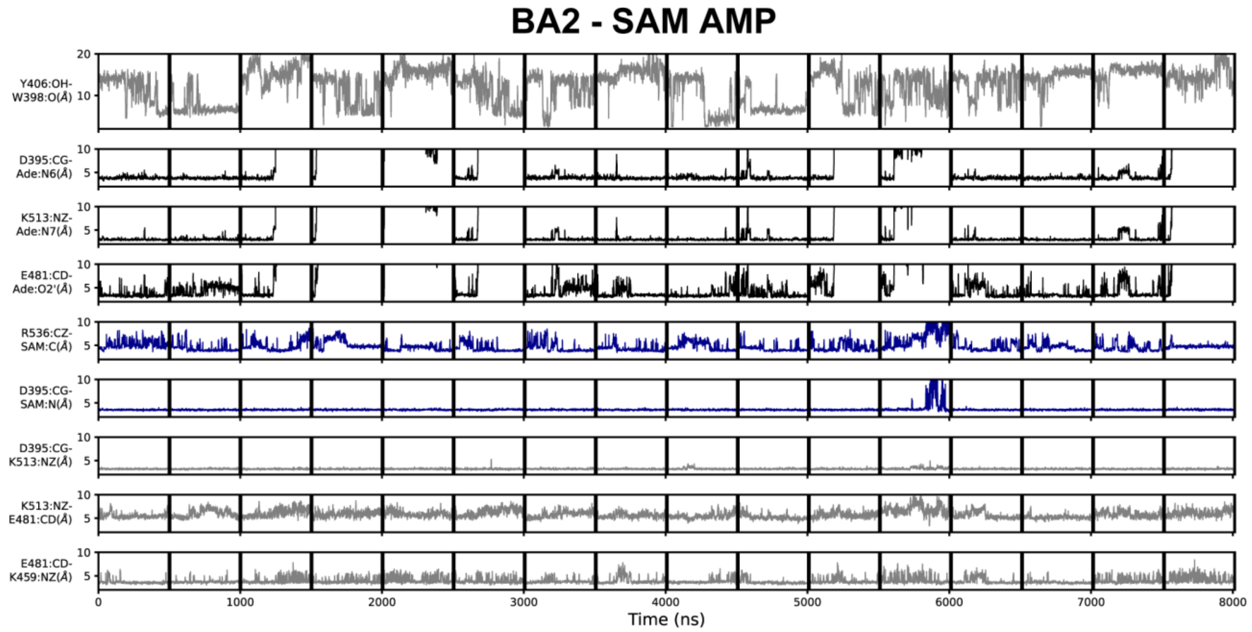


**Figure S8. Geometric annotation for trajectories started with substrates in the BA2 crystal structure conformation**. Shown are distance time series of 500-ns MD trajectories started from the BA2 conformation of METTL3 with (co)substrates. Starting conformation and bound ligands indicated on top of the Figure. Y406 to W398 distance, interaction of METTL3 to AMP substrate (black traces), METTL3 to SAM (blue traces), and intramolecular salt bridges.

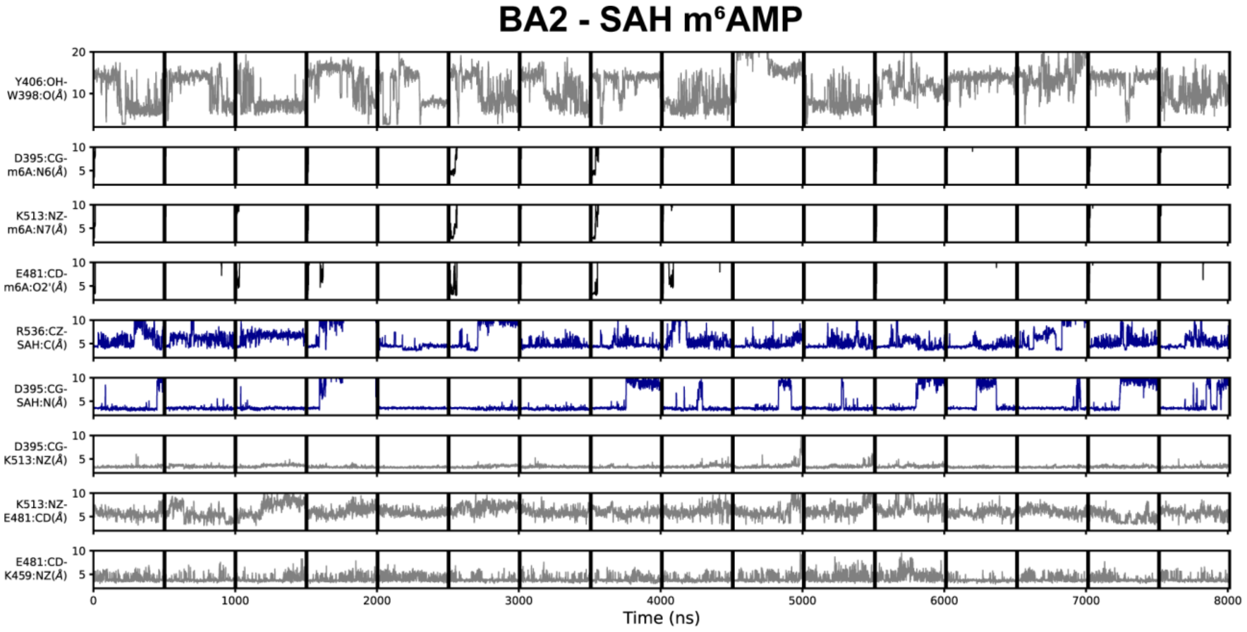


**Figure S9. Geometric annotation for trajectories started with substrates in the BA2 crystal structure conformation**. Shown are distance time series of 500-ns MD trajectories started from the BA2 conformation of METTL3 with (co)products. Starting conformation and bound ligands indicated on top of the Figure. Y406 to W398 distance, interaction of METTL3 to m$^6$AMP substrate (black traces), METTL3 to SAM (blue traces), and intramolecular salt bridges.

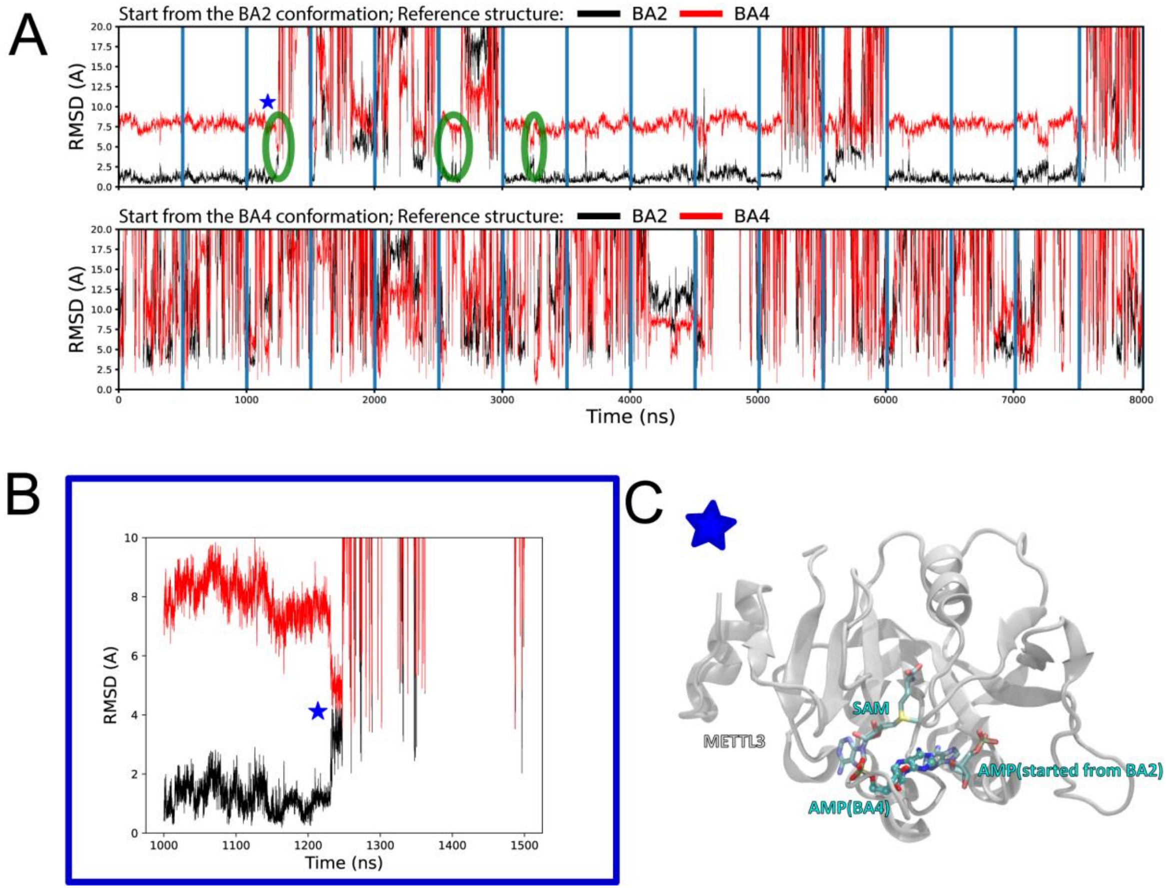


**Figure S10. Structural stability and conformational transitions of AMP in the MD simulations. (A)** Shown are time series of 500-ns MD trajectories of METTL3. The time series show the root mean square deviation (RMSD) of the adenine atoms of AMP started from the BA2 (top) or BA4 (bottom) conformation using as a reference their corresponding positions in the crystal structure of BA2 (black traces) or BA4 (red traces). AMP shows less RMSD fluctuations and is hence more stable in the MD simulations started from the complex in the BA2 (top) than BA4 (bottom) conformation. In some of the dissociation events from the BA2 conformation, AMP transiently populates a binding mode similar to the BA4 conformation (green circled regions and blue star). **(B)** Close up view of a trajectory starting from the BA2 conformation shows a dissociation event from the BA2 conformation in which AMP transiently populates a binding mode similar to the BA4 conformation (indicated with a blue star and green circle in (A)). **(C)** Conformation of the AMP at the frame marked by a star in (A) and (B) shows the AMP conformation overlaps with the one of AMP in the BA4 conformation (ball and sticks).

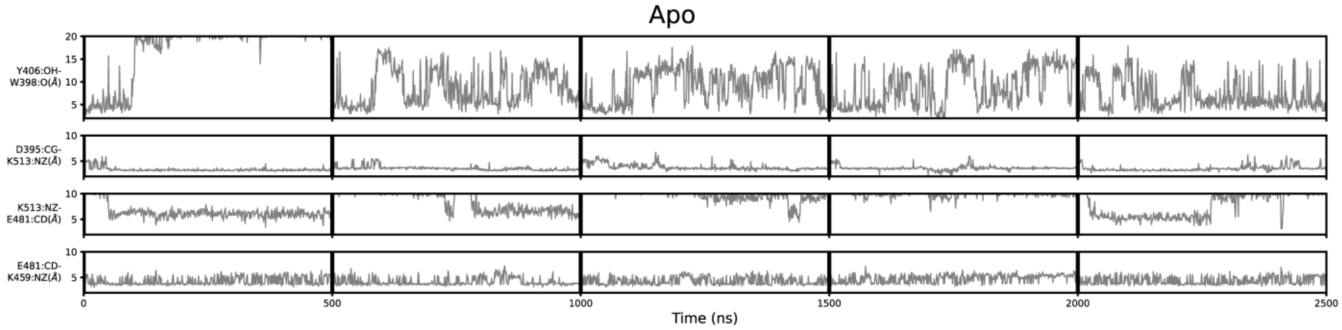


**Figure S11**. **Geometric annotation of apo trajectories**. Shown are distance time series of 500-ns MD trajectories of apo METTL3. Y406 sidechain in the ASL1 loop and intramolecular salt bridges monitored for five trajectories of apo METTL3-14.

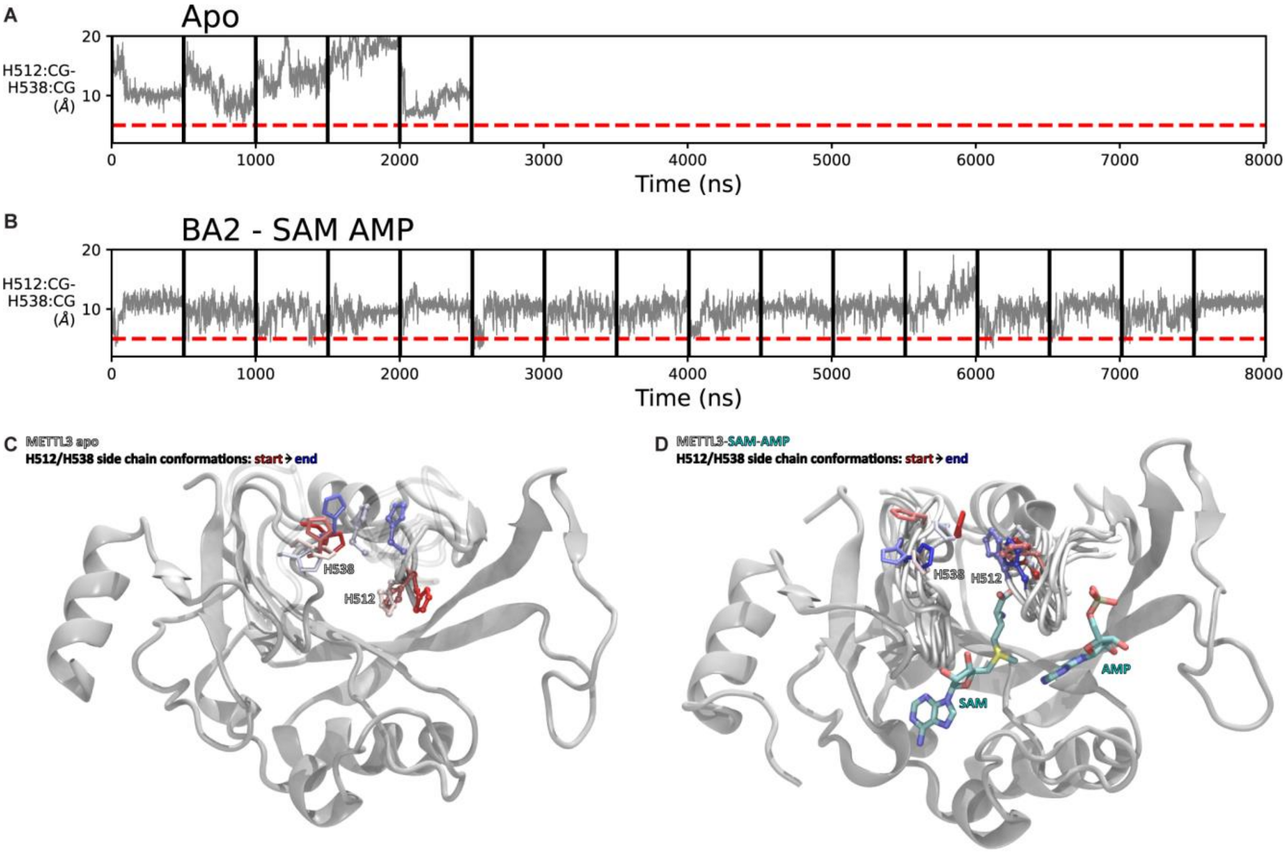


**Figure S12. Flexibility of the side chains of H512 and H538.** (**A,B**) Shown are distance time series between the Cγ atoms of H512 and H538 of 500-ns MD trajectories of apo METTL3 (A) and METTL3 bound to SAM and AMP that were started from the conformation of BA2 (B). (**C,D**) Conformation of H512 (ball and sticks) and H538 (sticks) at different time points (from red to blue) of a trajectory of apo METTL3-14 (C) and with bound SAM and AMP (D). In the presence of SAM, the motion of H512 is restricted. METTL14 was present in the MD simulation runs but is omitted here for clarity.

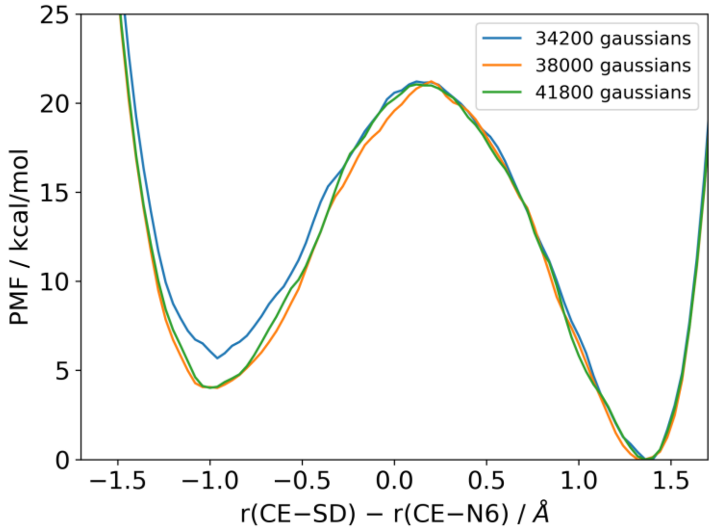


**Figure S13.** Potentials of mean force computed with different numbers of deposited Gaussians during the metadynamics simulation are compared to illustrate the convergence behavior of the simulation.

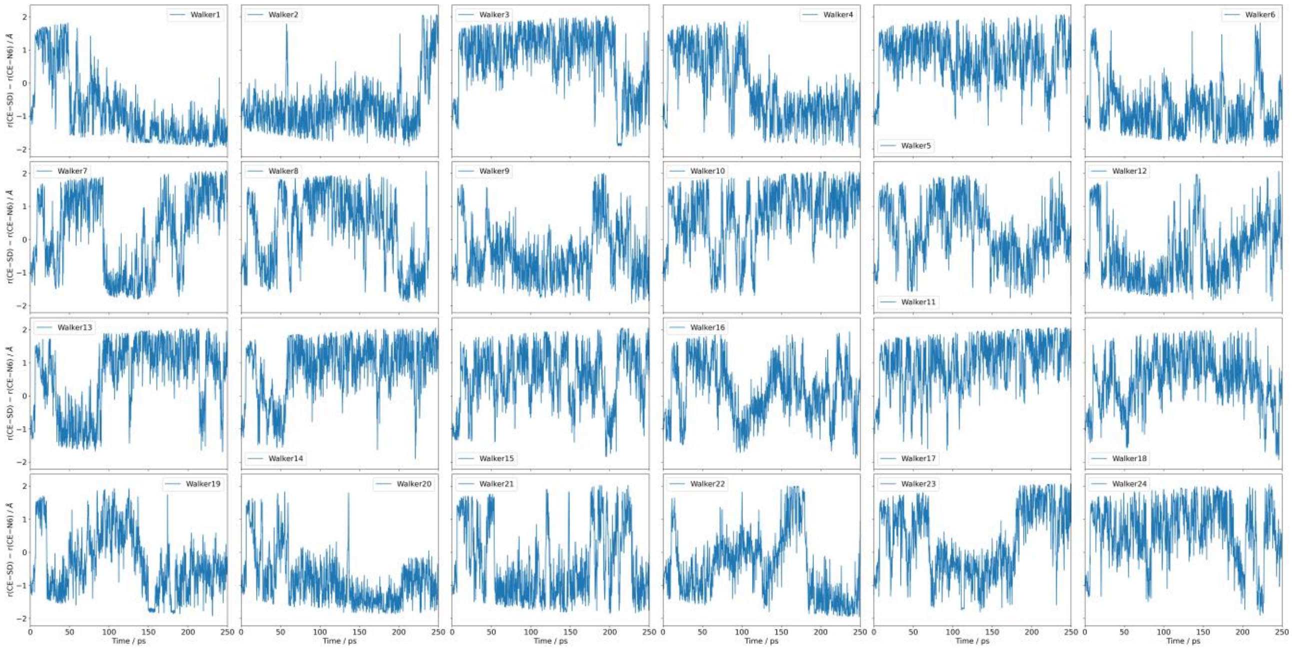


**Figure S14.** The time series of the collective variable (CV) sampled by the 24 independent walkers during one set of metadynamics simulations. These results clearly indicate that the CV exhibits diffusive behaviors between the reactant and product regions, further supporting the adequate sampling and convergence of the metadynamics simulations.

**Table S1.** Crystallography data collection and refinement statistics. Statistics for the highest-resolution shell are shown in parentheses.

| | BA1 complex (PDB 8PW9) | BA2 complex (PDB 8PW8) | BA4 complex (PDB 8PWA) | BA6 complex (PDB 8PWB) |
|---|---|---|---|---|
| **Data collection** | | | | |
| Wavelength (Å) | 1 | 1 | 1 | 1 |
| Resolution range (Å) | 49.69 - 2.301 (2.383 - 2.301) | 44.51 - 2.3 (2.382 - 2.3) | 44.54 - 2.1 (2.175 - 2.1) | 49.52 - 2.5 (2.589 - 2.5) |
| Space group | P 32 2 1 | P 32 2 1 | P 32 2 1 | P 32 2 1 |
| Unit cell (Å, °) | 63.94 63.94 225.14, 90 90 120 | 63.78 63.78 225.45, 90 90 120 | 63.89 63.89 225.18, 90 90 120 | 63.69 63.69 224.77, 90 90 120 |
| Total reflections | 237414 (22942) | 218643 (19326) | 311563 (28191) | 182166 (17587) |
| Unique reflections | 24644 (2396) | 24575 (2365) | 32185 (3137) | 19170 (1859) |
| Multiplicity | 9.6 (9.6) | 8.9 (8.1) | 9.7 (8.9) | 9.5 (9.4) |
| Completeness (%) | 99.80 (99.30) | 99.72 (99.41) | 99.72 (99.08) | 99.78 (99.79) |
| Mean I/sigma(I) | 17.40 (1.82) | 15.83 (1.61) | 18.28 (1.50) | 15.90 (1.75) |
| Wilson B-factor | 45.64 | 43.17 | 38.9 | 49.72 |
| R-merge | 0.101 (1.204) | 0.1072 (1.267) | 0.0969 (1.317) | 0.1241 (1.244) |
| R-meas | 0.1068 (1.272) | 0.1138 (1.351) | 0.1024 (1.398) | 0.1313 (1.315) |
| R-pim | 0.03422 (0.4072) | 0.0376 (0.456) | 0.03266 (0.4639) | 0.04229 (0.4233) |
| CC1/2 | 0.999 (0.644) | 0.999 (0.6) | 0.999 (0.61) | 0.998 (0.627) |
| CC* | 1 (0.885) | 1 (0.866) | 1 (0.87) | 1 (0.878) |
| **Refinement** | | | | |
| Reflections used in refinement | 24616 (2396) | 24523 (2365) | 32118 (3137) | 19134 (1859) |
| Reflections used for R-free | 1230 (120) | 1229 (119) | 1606 (157) | 958 (93) |
| R-work | 0.1921 (0.2542) | 0.1971 (0.2973) | 0.1907 (0.2731) | 0.1999 (0.3001) |
| R-free | 0.2383 (0.3146) | 0.2487 (0.3502) | 0.2250 (0.3064) | 0.2450 (0.3605) |
| CC(work) | 0.957 (0.834) | 0.956 (0.774) | 0.963 (0.814) | 0.954 (0.758) |
| CC(free) | 0.943 (0.675) | 0.949 (0.706) | 0.937 (0.809) | 0.937 (0.675) |
| Number of non-hydrogen atoms | 3648 | 3621 | 3742 | 3518 |
| macromolecules | 3473 | 3437 | 3480 | 3416 |
| ligands | 45 | 51 | 44 | 42 |
| solvent | 130 | 133 | 218 | 60 |
| Protein residues | 442 | 437 | 440 | 433 |
| RMS bonds (Å) | 0.019 | 0.018 | 0.018 | 0.019 |
| RMS angles (°) | 1.1 | 1.12 | 1.07 | 1.12 |
| Ramachandran favoured (%) | 96.73 | 96.91 | 97.89 | 95.68 |
| Ramachandran allowed (%) | 3.04 | 2.38 | 1.64 | 4.32 |
| Ramachandran outliers (%) | 0.23 | 0.71 | 0.47 | 0 |
| Rotamer outliers (%) | 0.28 | 0.28 | 0.55 | 1.69 |
| Clashscore | 3.38 | 3.85 | 3.65 | 6.71 |
| Average B-factor | 49.58 | 45.56 | 43.61 | 51.29 |
| macromolecules | 49.15 | 45.36 | 43.28 | 51.08 |
| ligands | 86.54 | 61.52 | 54.49 | 66.93 |
| solvent | 48.18 | 44.49 | 46.55 | 52.27 |

**Movie S1. Stable/flexible binding of SAM/AMP and flexibility of ASL1 loop/Y406 is evidenced by MD simulations.** Shown is a movie from MD simulations with the complex of METTL3 (grey ribbon representation) and METTL14 (teal ribbon representation) in complex with SAM (sticks representation, coloured by atom with carbon in teal and the sulphur atom in yellow) and AMP (sticks representation, coloured by atom with carbon in teal) started from the conformation of bisubstrate analogue BA2. METTL3 residues D395, W398, Y406, K459, E481, S511, K513, and R536 are shown as sticks and coloured by atom with carbon in teal. The movie begins with the SAM and AMP in the position of BA2. The flexibility of the Y406 sidechain and the ASL1 loop in which it is located is observed already at the beginning of the movie. At around time point 00:15, the Y406 interacts with the adenosine ring of AMP, and at 00:20, there is a dissociation of AMP from the binding pocket. AMP remains bound to Y406 via $\pi$-$\pi$ interactions for the next 10 s and finally dissociates completely. In the natural system, the adenosine would remain bound by the rest of the RNA chain, but in this case, when AMP is alone, it dissociates away. Y406 occupies the binding pocket close to SAM. At around 00:39, the AMP re-enters in the vicinity of the pocket, and the Y406 again coordinates via $\pi$-$\pi$ interactions with the adenosine ring at 00:45. After around 10 s, the AMP dissociates again. The binding mechanism of the natural substrate probably requires the binding of the RNA chain in the groove, followed by the interaction of Y406 with the substrate adenosine. The rest of the movie continues with a partial dissociation of the methionine moiety of SAM at around 01:05 and the non-specific interactions of AMP with METTL3-14.
